# PDAUG - a Galaxy based toolset for peptide library analysis, visualization, and machine learning modeling

**DOI:** 10.1101/2021.02.02.429203

**Authors:** Jayadev Joshi, Daniel Blankenberg

**Affiliations:** Genomic Medicine Institute, Lerner Research Institute, Cleveland Clinic, Cleveland, Ohio, USA; Department of Molecular Medicine, Cleveland Clinic Lerner College of Medicine, Case Western Reserve University, Cleveland, Ohio, USA

## Abstract

Computational methods based on initial screening and prediction of peptides for desired functions have been proven effective alternatives to the lengthy and expensive methods traditionally utilized in peptide research, thus saving time and effort. However, for many researchers, the lack of expertise in utilizing programming libraries and the lack of access to computational resources and flexible pipelines are big hurdles to adopting these advanced methods. To address these barriers, we have implemented the **P**eptide **D**esign and **A**nalysis **U**nder **G**alaxy (PDAUG) package, a Galaxy based python powered collection of tools, workflows, and datasets for a rapid *in-silico* peptide library analysis. PDAUG offers tools for peptide library generation, data visualization, in-built and public database based peptide sequence retrieval, peptide feature calculation, and machine learning modeling. In contrast to the existing methods like standard programming libraries or rigid web-based tools, PDAUG offers a GUI based toolset thus providing flexibility to build and distribute reproducible pipelines and workflows without programming expertise. Additionally, this toolset facilitates researchers to combine PDAUG with hundreds of compatible existing Galaxy tools for limitless analytic strategies. Finally, we demonstrate the usability of PDAUG on predicting anticancer properties of peptides using four different feature sets and assess the suitability of various machine learning algorithms.

## Introduction

The interest in peptides related research has been gaining in popularity over the last decades (Lee et al., 2019). A large number of naturally occurring peptides (over 7,000) with potentially important roles in human physiology have been identified and, currently, more than 140 peptide therapeutics are in different stages of clinical trials (Fosgerau and Hoffmann, 2015). In view of their integral importance in a number of signal transduction pathways, they are ideal candidates for functioning as drugs, especially as anticancer or antimicrobial peptides (Adermann et al., 2004). Usually, peptides are naturally occurring molecules synthesized by cellular processes and which adopt alternative conformations according to their biological functions (Lee et al., 2019). They can act as natural ligands in the form of cofactors, coenzymes, hormones, or directly interact with macromolecules like protein, RNA, or DNA (de Araujo et al., 2019). The research underlying the design of therapeutic peptides, such as peptide based drugs and vaccines, demands intense effort and assets in establishing their pharmacokinetic and pharmacodynamic properties such as serum stability, bioavailability toxicity, etc. (Bray, 2003; Otvos and Wade, 2014). Peptide-based vaccine approaches have emerged as a powerful candidate in many therapeutic areas, including infectious diseases and cancer (Malonis et al., 2020) Characterization of peptides that bind to specific MHC molecules is therefore of great importance for peptide-based vaccines. However, in comparison to expensive and lengthy biochemical experiments, bioinformatics methods for predicting MHC binding peptides have been very popular in recent times (Kalita et al., 2020; Oyston and Robinson, 2012; Sidro-Llobet et al., 2019). Various computational approaches have been shown to offer the best cost-benefit ratio across translational research areas (Saeb, 2018; Wu et al., 2012; Xia, 2017). Leveraging *in-silico* approaches to uncover peptides with desired pharmacological action can be expected to significantly lower the cost and time required to establish a drug or a vaccine candidate (Lavecchia and Di Giovanni, 2013). In fact, computational prediction of peptides with desired functions have been providing effective alternatives to the lengthy and expensive traditional methods in peptide research thus saving time and efforts (Bhadra et al., 2018; Hamid and Friedberg, 2019; Jabbar et al., 2018; Lata et al., 2010; Meher et al., 2017; Schaduangrat et al., 2019). The concept of prioritizing sequence-based properties of a protein sequence as a function of sequence-derived features is not new (Karlin and Altschul, 1990). Over the past decade, approaches based on physicochemical, compositional properties, k-mer counting, etc. have been proposed (Chou, 2001; Saidi et al., 2010; Yao et al., 2019). With the rise of computational power, feature-based methods evolved profoundly, expanding into the 3D structure level of biomolecules (Hicks et al., 2019). However, necessary programming and mathematics expertise, as well as limitations in hardware resources are the core challenges associated in utilizing programming based resources (Kesh and Raghupathi, 2004; Rhee, 2005). On the other hand, existing web-based tools do not provide pipelines that are both reproducible and flexible, failing to solve the major hurdles in adopting these methods in research (Gilbert, 2004). Platforms, such as Galaxy (Afgan et al., 2018; Giardine et al., 2005), have emerged to address the needs of researchers in a user-friendly manner. Galaxy is an open, web-based platform for accessible, reproducible, and transparent computational research, providing a wealth of computational tools and visualizations, analysis workflows, and training materials.

Here, we present PDAUG, a Galaxy tool suite that includes 24 different tools for the analysis of peptide libraries. With this toolset, we have provided a set of functionalities that includes peptide library generation, feature analysis, data visualization and plotting, machine learning modeling, and dataset retrieval in a user-friendly manner under one platform. These modular command-line tools leverage Galaxy to provide an interactive graphical interface for each individual tool as well as an expandable set of workflows for peptide data representation and analysis. Individual tools rely on pandas dataframes to handle the data matrices, with tabular and FASTA formats for input/output (IO) operations. Data formats were chosen for PDAUG to complement the strengths of Galaxy’s existing toolsets and to support user expectations. Tests have been defined for each tool to maintain reliable and reproducible results. In addition, we have produced an interactive Galaxy tutorial for each example workflow used in this article, demonstrating the functionality and usability of this toolset. Finally, we utilized this toolset to assess a suitable combination of features and machine learning algorithm in predicting the anti-cancer properties of peptides to demonstrate the usability of this toolset in the peptide research.

### Implementation

A graphical overview of PDAUG has been described in Figure 1. A detailed description of each tool can be found in Table 1.

**Table 1.** Description PDAUG tools.

| Functionality | Tool name | Major Libraries used |
| --- | --- | --- |
| Data visualization and plotting | PDAUG Basic Plots | matplotlib*, pandas*, seaborn*, quantiprot |
|  | PDAUG Fishers Plot | PDAUG Fishers Plot |
|  | PDAUG Peptide Data Plotting |  |
|  | PDAUG Peptide Ngrams |  |
|  | PDAUG Sequence Network |  |
|  | PDAUG Peptide Length Distribution |  |
|  | PDAUG Uversky Plot |  |
| Descriptor calculation | PDAUG AA Property Based Peptide Descriptor | modlAMP, pandas, pydpi |
|  | PDAUG Peptide Core Descriptors |  |
|  | PDAUG Peptide Global Descriptors |  |
|  | PDAUG Sequence Property Based Descriptors |  |
|  | PDAUG Word Vector Descriptor |  |
| Peptide library generation | PDAUG AA Property Based Peptide Generation | modlAMP, pandas |
|  | PDAUG Sequence Based Peptide Generation |  |
| Machine learning | PDAUG ML Models | sklearn*, matplotlib, seabornpandas, gensim, nltk |
|  | PDAUG Word Vector Model |  |
| Circular dichroism (CD) data analysis | PDAUG Peptide CD Spectral Analysis | modlAMP, pandas |
| Peptide 3D structure | PDAUG Peptide Structure Builder | fragbuilder, pandas |
| Core functionality | PDAUG Peptide Sequence Analysis | modlAMP, pandas |
|  | PDAUG Peptide Core Functions |  |
| Peptide data access | PDAUG Peptide Data Access | modlAMP, biopython, pandas. |
| Data handling & IO | PDAUG TSVtoFASTA | pandas |
|  | PDAUG Merge DataFrames |  |
|  | PDAUG AddClassLabel |  |
| *Python libraries used in data science. |  |  |

**Figure 1.**
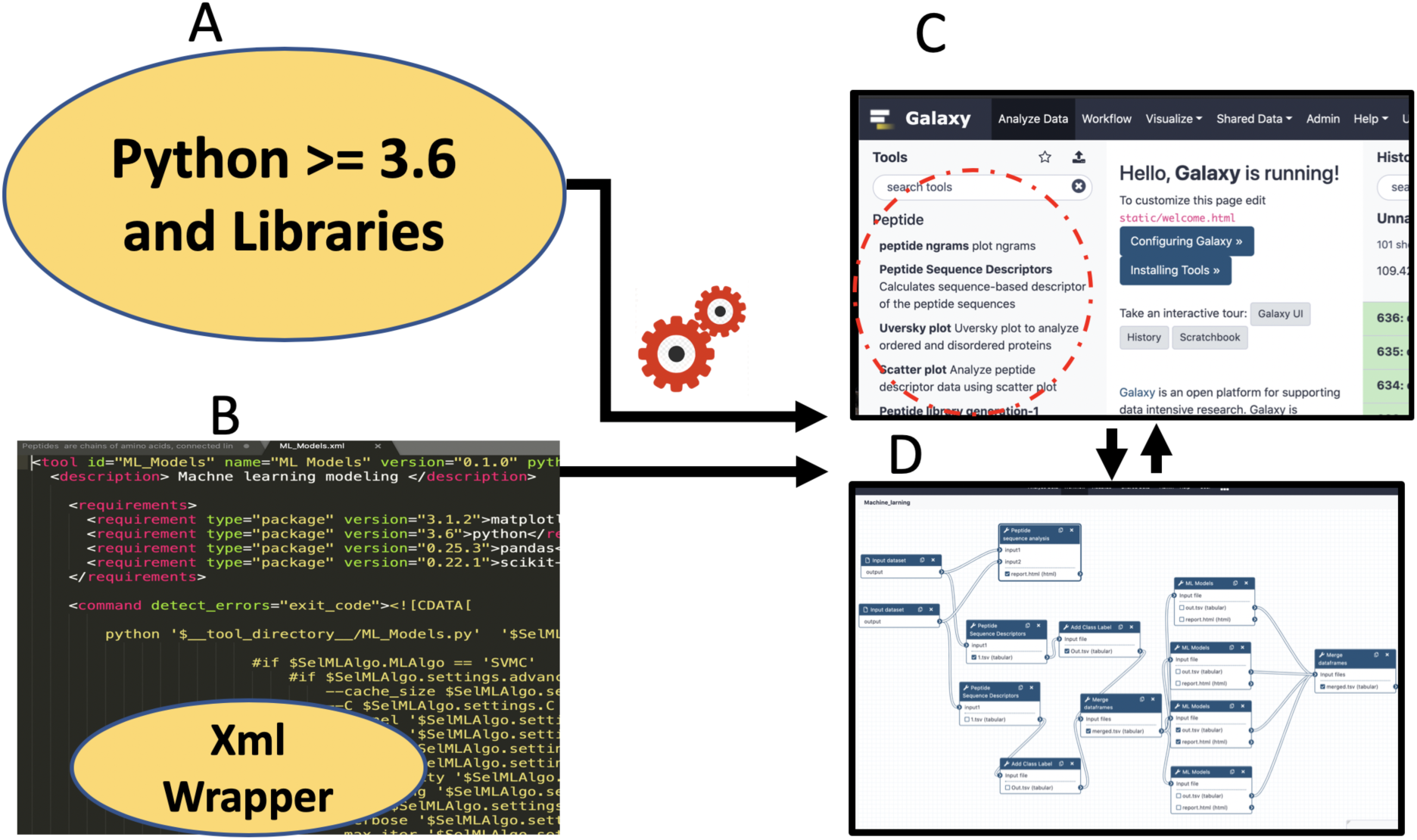
Extending peptide library analysis with the PDAUG toolset inside Galaxy. **(A)** Tools created with python libraries. (B) Implementing Galaxy tool wrappers and tests for each tool. (C) PDAUG tool set with 24 individual tools. (D) Implementing reusable workflows using PDAUG.

### Programming languages

Due to its popularity among the scientific community, we have chosen to implement the functions and backend scripts for Galaxy tools in Python. We have leveraged popular scientific libraries such as NumPy, SciPy, pandas, Matplotlib, scikit-learn (sklearn), etc. for data manipulation and representation to maintain uniformity and simplicity. Galaxy tool wrappers have been designed and uploaded to the ToolShed (Blankenberg et al., 2014), enabling point-and-click installation. We also provide a Docker image with Galaxy and the tools pre-installed (https://hub.docker.com/r/jayadevjoshi12/galaxy_pdaug; https://github.com/jaidevjoshi83/docker_pdaug).

### Accessing peptide data from pre-populated local and remote web based resources

In addition to allowing the upload of user-provided datasets, PDAUG has been equipped with “PDAUG Peptide Data Access” tool for quick and easy access to various publicly available peptide datasets. This tool implemented based on modlAMP (Müller et al., 2017) and Biopython (Cock et al., 2009) and includes antimicrobial peptides (AMPs), trans-membrane peptides, peptides from the UniProt database, anticancer peptides (ACPs), helical transmembrane peptides (HTPs) and randomly scrambled AMPs. Additionally, we have also provided option to fetch data directly from the two popular web resources: Antimicrobial Peptide Database (APD) (Wang et al., 2016) and Collection of Antimicrobial Peptides Database (CAMP) (Waghu et al., 2016).

### Peptide library generation

Two tools with several options to generate peptide sequences have been implemented. These tools provide various methods based on amino acid and sequence properties to generate peptide sequence libraries with different properties which can be utilized to further analysis inside Galaxy.

1. **PDAUG AA Property Based Peptide Generation**. This tool generates sequences mostly based on amino acid (AA) properties. The user can generate peptide sequences based on 10 different options which include, option “AmphipathicArc Peptides’’ returns peptides with presumed amphipathic helices, “AMPngrams Peptides “ returns peptides from the most frequent n-grams in the APD3 database. “Centrosymmetric Peptides’’ returns centrosymmetric peptides sequences with a symmetry axis, “Helices Peptides” returns presumed amphipathic helical peptides, “HelicesACP Peptides” returns peptides with the AA probability similar to helical ACPs, “Kinked Peptides’’ returns peptides with presumed amphipathic helices with a kink, “Hepahelices Peptides” returns peptide with presumed amphipathic helices and a heparin-binding-domain, “Oblique Peptides” returns presumed oblique oriented peptides, “Random Peptide”, returns random peptides with a specified amino acid distribution, and “MixedLibrary Peptides “ returns a library of mixed peptides. Most of the functions of this tool have been implemented on the top of modlAMP and pandas libraries.
2. **PDAUG Sequence Based Peptide Generation**. This tool generates peptide libraries based on three different options. The primary method “Random Peptides” is based on permutation and combinations that perform a search for all the possible combinations of 20 amino acids within the given length. The second method “Mutated Peptides” produces replacement of existing amino acids with the remaining 19 possibilities at given positions. The last method “Sliding Window Peptides” takes an input of a protein sequence and generates random peptide fragments based on a sliding window and fragment size.

### Peptide Structure

A peptide sequence can be folded and adopts a unique structural space for specific biological functions. The tool “PDAUG Peptide Structure Builder” has been implemented to generate a peptide structure based on the libraries FragBuilder (Christensen et al. 2014) and Open Babel (O’Boyle et al. 2011). This tool can generate peptide sequences up to 4 amino acids that can be utilized in small peptide docking simulations with molecular docking tools such as AutoDock Vina (Trott and Olson, 2010).

### Peptide descriptor generation

There are four different tools that have been implemented that calculate more than 10,000 descriptors based on 50 different classes of peptide descriptors for a given peptide sequence. We have implemented four different tools to calculate peptide descriptors.

1. **PDAUG Peptide Global Descriptors**. This tool calculates simple one-dimensional peptide descriptors based on 11 different options which include Sequence Length, Molecular Weight, Sequence Charge, Charge Density, Isoelectric Point, Instability Index, Aromaticity, Aliphatic Index, Boman Index and All. These descriptors are important to define the global properties of a peptide sequence and can be utilized to build machine learning models to predict a biological property.
2. **PDAUG Sequence Property Based Descriptors**. This tool calculates descriptors based on 13 different options. Option “GetAAComp” calculates amino acid composition descriptors, “GetDPComp” calculates dipeptide composition descriptors “GetTPComp” calculates tri-peptide composition descriptors, “GetMoreauBrotoAuto” calculates normalized Moreau-Broto autocorrelation descriptors, “GetMoranAuto calculates” moran autocorrelation descriptors, “GetGearyAuto” calculates Geary autocorrelation descriptors, “GetCTD” calculates composition Transition Distribution descriptors, “GetPAAC” calculates Type I Pseudo amino acid composition descriptors, calculates “GetAPAAC” amphiphilic (Type II) Pseudo amino acid composition descriptors, “GetSOCN” calculates sequence order coupling numbers, “GetQSO” calculates quasi sequence order descriptors, “GetTriad” calculates the conjoint triad features from the protein sequence and, “BinaryDescriptor” calculates the binary descriptor of peptides with identical lengths. Lastly, the “All” option calculates all the above descriptors with one click, except the binary descriptors. These descriptors are implemented based on PyDPI library (Cao et al., 2013).
3. **PDAUG AA Property Based Peptide Descriptor**. This tool calculates descriptors derived from amino acid properties based on six different options. “Calculate AutoCor” computes descriptors via auto-correlating the amino acid values. “Calculate CrosCor” computes descriptors via cross-correlating the amino acid values. “Calculate Movement” computes a descriptor based on the maximum or mean movement of the amino acid values. “Calculate Global” option computes descriptors via calculating a global / window averaging descriptor values. “Calculate Profile” computes descriptors via calculating hydrophobicity or hydrophobic moment profiles for given sequences and fitting for slope and intercept. “Calculate Arc” computes descriptors via calculating property arcs. These descriptors depend upon the given descriptor scale and window size.
4. **PDAUG Word Vector Descriptor**. Word2vec is a popular technique of word embedding in which words from a vocabulary are represented as vectors based on similar contexts. The similarity of words derived from proximity, which is calculated based on a large corpus of documents. (Mikolov *et al*., 2013; Asgari and Mofrad, 2015). Such representation of protein sequence data has led to better performance in protein and peptide classification (Hamid and Friedberg, 2019; Wu et al., 2019; Yang et al., 2018). In this toolset, we have included two tools: the first tool “PDAUG Word Vector Model” generates a word2vec model that contains the contextual information for each trigram in the corpus of given protein sequences. Input protein sequences referred to as corpus utilized to generate trigram-based vocabulary. Gensim library (Rek and Sojka, 2010) is used to apply a continuous bag of words (CBOW) or skip-gram algorithm to generate a 200-dimensional vector for each trigram. These 200-dimensional vectors represent the context information of all the trigrams present in the training. These vectors can be utilized to generate the descriptor for the peptides using the second tool “PDAUG Word Vector Descriptor”. A precalculated skip-gram word2vec model, generated based on UniProtKB/TrEMBL database (Hamid and Friedberg, 2019), has been provided with the supplementary data as model.txt, which can be utilized directly with the “PDAUG Word Vector Descriptor” tool to calculate 200 descriptors.

### Data visualization and analysis

PDAUG contains several data visualization tools for both sequence and feature based data representations.

1. **PDAUG Basic Plots**. This tool is equipped with four different options to plot the data in the tabular and FASTA format. Option “Heat Map” plots heatmap for the given data set. “Box Plot” generates the box plot for given data based on the class label. “Scatter Plot” generates 2D or 3D scatter plots for 2 or 3 features and highlights data based on the class labels. Each of these three plotting methods uses tabular data. Option “Word Cloud” has been implemented to graphically assess a set of peptide sequences on the basis of n-grams. Word clouds are commonly used to graphically represent various n-grams frequencies in a particular population.
2. **PDAUG Fisher’s Plot**. Fisher plot has been implemented to assess two peptide sequences on the basis of their feature spaces. In principle, the Fisher’s plot compares two peptide sequences in two-dimensional spaces, defined by quantitative features of peptide sequences. This tool computes Fisher’s exact test on a local and global ratio of peptide sequence in feature space where the global and local ratio is computed either in the whole feature space or in feature space belonging to each set. This tool is implemented based on the Quantiprot (Konopka et al., 2017) package and can be utilized to compare two peptide libraries.
3. **PDAUG Peptide Data Plotting** This tool provides four different plotting options. “Helical Wheel” plots a helical wheel plot for a given peptide sequence. “Probability Density Estimation” plots probability density estimations of given data vectors. “Violin Plot” creates a violin plot from the given data array. “Amino Acid Distribution” plots the amino acid distribution of a given sequence library.
4. **PDAUG Peptide Ngrams** Distribution of n-grams varies from sequence to sequence which affects the property of sequences. This tool counts n-grams in the entire peptide sequence data and fits their distribution with the Zipf’s law, also known as power-law distribution.
5. **PDAUG Sequence Similarity Network** This tool calculates Levenshtein distance between peptide sequences, and plots the data in the form of a sequence similarity network. A dispersed and highly clustered network represents less similarity between sequences. Conversely, a network that is compact and has a smaller number of clusters represents high sequence similarity between sequences.
6. **PDAUG Uversky Plot**. Uversky plot was first proposed by Uversky (Uversky, 2002) to understand the ordered and disordered protein sequences. Uversky plot separates proteins into globular and intrinsically disordered protein subsets on the basis of their mean net charge versus mean hydropathy. Uversky plot has been implemented under the tool name “PDAUG Uversky Plot”, where users can compare two different peptide libraries on the basis of their globular and intrinsically disordered properties (Konopka et al., 2017).
7. **Summary Plot** Summary plot options of the “PDAUG Peptide Sequence Analysis” tool consists of six subplots. Subplots include a) bar graph that compares two peptide libraries on the basis of their AA fractions, b) bar graph that compares two libraries on the basis of global charge fraction, c) box plot that compares two libraries on the basis of sequence length distribution, d) violin plot that compares two libraries on the basis of global hydrophobicity, e) violin plot that compares two libraries on the basis of global hydrophobic movement, and f) 3D scatter plot that compare two libraries on the basis of their global hydrophobicity, global hydrophobic movement, and global charge.
8. **PDAUG Peptide Core Functions**. This tool is equipped with four options, which includes “Mutate Amino Acids” mutates a number of positions per sequence randomly with a given probability value, “Filter Duplicates” removes duplicate sequences from a library, “Keep NaturalAA” filters out sequences with unnatural amino acids, and “Filter Amino Acids” filters out sequences with specific amino acids.
9. **PDAUG Peptide Sequence Analysis**. This tool provides utility to calculate a few important sequence-based properties and analyze peptide library sequences based on these properties. With this tool six different options have been provided which includes, option “Calculate Amino Acid Frequency” calculates amino acid frequency in the given peptide library, option “Calculate Global Hydrophobicity” calculates global hydrophobicity for all sequences in the library, option “Calculating Hydrophobic Moments” calculates hydrophobic moments for all sequences in the given peptide library, option “Calculate the total molecular charge” calculates the total molecular charge at a given pH for all sequences in the library, option “Calculate the sequence length” calculates sequence length of all sequences in the library and option “Summary Plot” compares two peptide libraries based on above properties and generates a visual summary.

### Machine learning model building, cross-validation, and accuracy assessment

PDAUG provides standard utilities for machine learning modeling and model selection via the “PDAUG ML Models” tools. These tools can classify peptides in a binary fashion and predict different peptide classes. A total number of seven different supervised and one artificial neural network algorithms, including Logistic Regression classifier (LRC) (Stoltzfus, 2011), Gaussian Naive-Bayes classifier (GNBC) (Friedman et al., 1997), K-nearest neighbor classifier (KNBC) (Cunningham at el. 2007), Decision-tree classifier (DTC) (Jenhani et al., 2008), Support vector machines classifier (SVMC) (Cortes et al. 1995) Random Forest classifier (RFC) (Liaw et al. 2002), Gradient boosting classifier (GBC) (Natekin and Knoll, 2013), Stochastic gradient descent classifier (SGDC) (Zhang, 04), and multilayer perceptron (MLP) (Pal and Mitra, 1992), have been implemented. Cross-validation has been included in our methodology for accuracy estimation (Kohavi, 1995). In cross-validation, active and inactive data were randomly divided into n-folds or parts (up to 10), where each set has the nth part of active as well as inactive peptides. The algorithm was trained on the n-1 sets and prediction was made on the remaining set. This process was repeated for each and every set. Thus, the final performance scores of accuracy, precision, recall, f1-score, and AUC were calculated as a mean for all the folds. Two normalization methods, max to min and z-scaling, to normalize the data before computational modeling is also implemented. To test the model, 4 different options have been made available. “Internal Test”, which uses cross validation to assess redundancy of model for testing the model. “Train-Test Split”, which splits training data into two fractions, one fraction as training data and another as for testing data. “External Test Data”, which enables the inclusion of a separate data file as a testing data set other than training data. And finally, “Predict unknown”, which can be used to make predictions on unknown datasets, have been implemented.

### Circular dichroism spectral data analysis

Secondary structure dynamics may be a major feature in determining various biological properties of different classes of peptides. Circular dichroism (CD) is a potential technique to determine the secondary structure and folding properties of proteins (Ranjbar and Gill, 2009). Initial laboratory experiments usually include CD spectroscopy of peptides in different solvents. We have included “PDAUG Peptide CD Spectral Analysis”, a tool based on the modlAMP and pandas libraries, that can be used to analyze CD spectroscopy of peptides data in different solvents. This tool handles CD data based on 4 different options. “Calculate Ellipticity” calculates molar ellipticity and mean residue ellipticity for all the tabular data provided. “Generate CD Plots” generates the CD plots. “Save Data in DichroWeb Readable Format” converts and returns data into DichroWeb compatible format. And “Calculate the Percent of Helicity”, which calculates percentage of helicity based on mean residue ellipticity data.

### IO operations

All the tools were implemented based on pandas data frame for seamless data operation between the various tools. We have included three tools to handle and manipulate FASTA and tabular data files. “PDAUG TSVtoFASTA” tool changes the input data formats from tabular to FASTA and splits the data on the basis of their class label in separate FASTA files. “PDAUG AddClassLabel”, add a desired class label to samples in a dataframe frame. The last tool of this category is “PDAUG Merge Dataframes” which merges two user-provided data frames. This simplifies IO operations allowing PDAUG to interact with already existing Galaxy tools.

### Example data

A high-quality dataset was extracted from a previously published work to demonstrate the usability of the described toolset (Hajisharifi et al., 2014). Initial data contains 192 anticancer peptides ACPs and 215 non-ACPs. In order to remove the redundancy and ensure the quality, the example dataset was cleaned as described in previously published work (Schaduangrat et al., 2019). Finally, a count-balanced dataset, 138 ACPs (positive) and 138 non-ACPs (negative) sequences was obtained by randomly removing sequences from the negative set. The length distribution of the positive dataset is somewhat different from the negative dataset, primarily due to the presence of several longer outlier sequences. Figure 2 depicts the length distribution of the positive and negative datasets. A sequence similarity network was calculated and presented in Figure 3, which shows less diversity in ACPs in comparison to non-ACPs.

**Figure 2.**
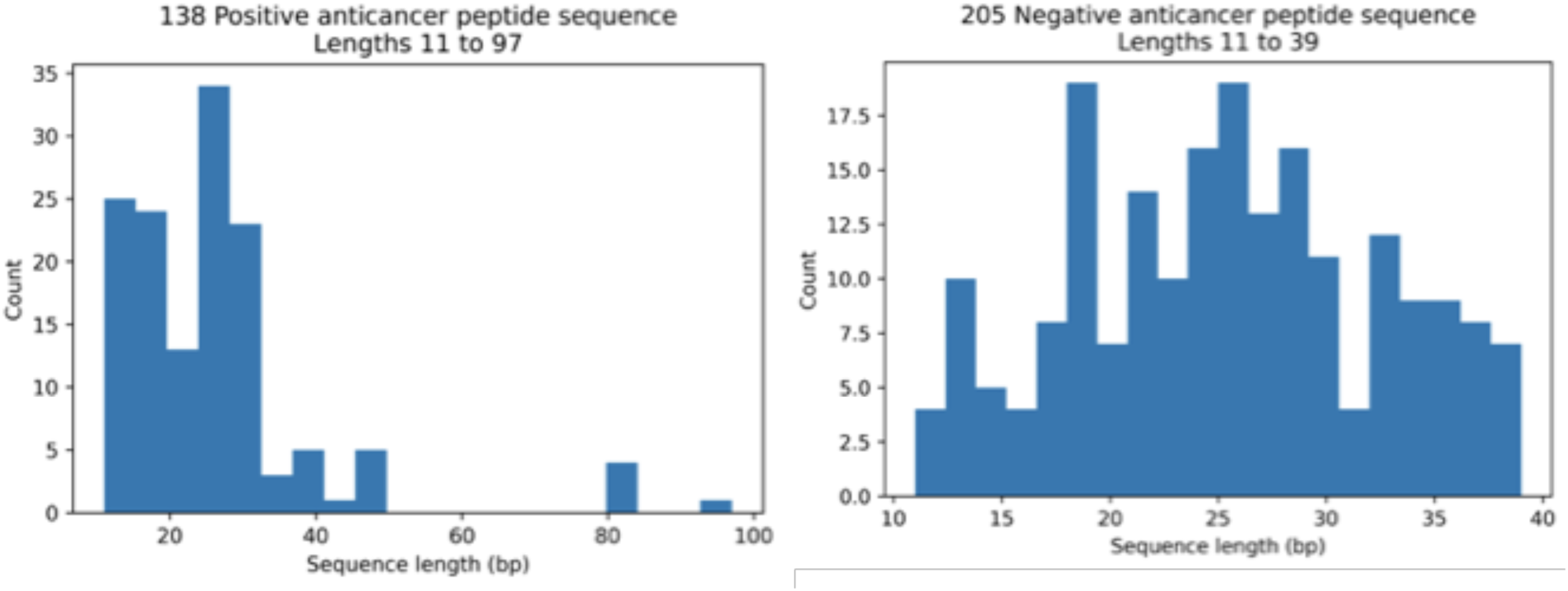
Sequence length distributions for the anticancer peptide and non-anticancer peptides. Mean lengths of anticancer and non-anticancer peptides are 40.06 and 32.25 amino acids, respectively, with less variability in length shown among the anticancer peptides.

**Figure 3.**
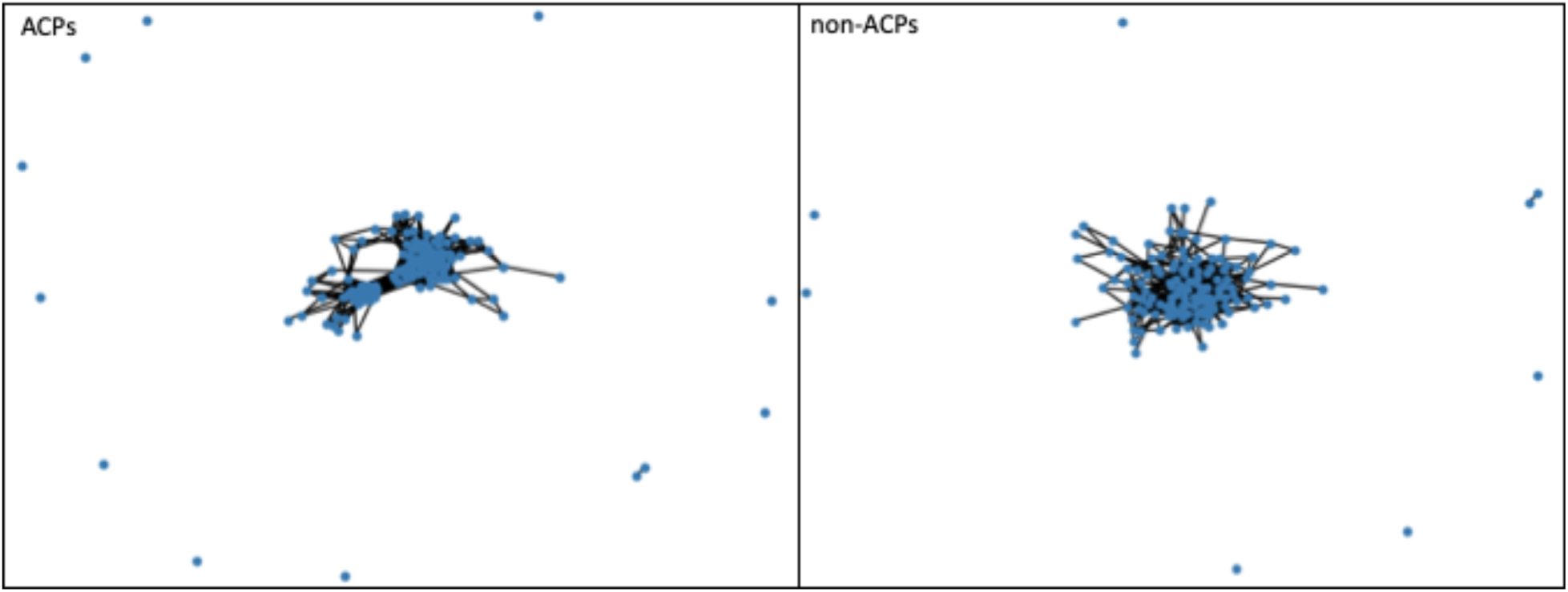
Sequence similarity network of the ACPs and non-ACPs. In comparison to the non-ACPs peptides, ACPs show two compact clusters that indicate a relatively high sequence similarity. In case of non-ACPs, relatively scattered networks have been observed.

### Descriptor Calculation

We have used four different descriptors. 1) Composition, Transition, Distribution descriptors (CTD), where Composition is expressed as the ratio of a number of amino acids of a particular property versus a total number of amino acids. Transition descriptors describe the relation between the transitions from a particular property to different properties and the number of amino acids. 2) Geary autocorrelation descriptors (GearyAuto), 3) Moran autocorrelation descriptors (MoranAuto), Moran, and Geary autocorrelation descriptor describes how a property is distributed along with the topology where special correlation can be represented through Moran or Geary correlation coefficient, and 4) Word Vector descriptor was calculated based on word embedding using skip-gram algorithm. A total number of 140, 200, 200 and 200 descriptors can be calculated for all these methods, CTD, GearyAuto, MoranAuto, and Word Vector, respectively.

### Machine learning workflow

Six algorithms, LRC, RFC, GBC, DTC, SGDC, and SVMC, have been applied to the training dataset and 10-fold cross-validation was used for accuracy estimation. In cross-validation, positive and negative data were randomly divided into 10 folds or parts, where each set has the 10th part of active as well as inactive peptides. The algorithm was trained on the 9 sets and the prediction was made on the remaining 10th set. This process was repeated for each and every set. Thus, the final performance scores were calculated as a mean of all the folds. We also included two methods, max to min and z-scaling, to normalize the data before ML modeling. The entire workflow was applied to the four descriptor sets and the performance was estimated based on routinely used performance measures: a) **Precision**, also known as the probability of positive values (PPV), is summarized as the probability of currently predicted positive instances and estimated on the basis of True Positive (TP) and False positive (FP). b) **Recall**, also known as sensitivity, is defined as the estimation of the percentage of the correctly predicted positive instances, and is also calculated with TP and FP. c) **F1** measures also an important estimate in model accuracy and can be defined as a harmonic mean of precision. The value for each of these three estimates falls between 0 and 1, with larger values indicating better performance and better accuracy. d) **Accuracy** is described as correctly predicted instances and calculated on the basis of TP and True Negative (TN) divided by TP, TN, FP, and False Negative (FN) e) AUC is Area under ROC curve, where ROC is a receiver operating characteristic. AUC represents the area covered by ROC.

## Result and discussion

To meet the increasing popularity of the design and screening of large peptide libraries, we have developed this toolset for peptide library analysis. Traditional peptide library design and analysis is labor-intensive work that requires bioinformatics methods to enable scalable alternatives. To maximize accessibility and impact, we have released PDAUG as a set of Galaxy tools, enabling web-based access and sharing of tools, pipelines, and analysis results. We have implemented 23 different tools for visualization, analysis, and library generation for peptides research.

### Anticancer / non-anticancer peptide dataset

A total number of 138 ACPs and 138 non-ACP sequences were selected for the final training data set. Figure 2 describes the length distribution of the ACP and non-ACP sequences. Mean lengths of ACPs and non-ACPs are observed somewhere in the range of 32 to 40 amino acids. However, fewer outlier sequences, more than 90 amino acids are observed in the ACP dataset. The sequence similarity network calculated by Levenshtein distance algorithms shows two compact clusters in the ACP dataset, conversely, a comparatively scattered network is observed in the case of the non-ACP dataset. The sequence similarity network shows relatively fewer diverse sequences in the ACP data set in comparison to the non-ACP sequence (Figure 3). A detailed summary plot calculated with the help of the “PDAUG Peptide Sequence Analysis” tool, compares ACP and non-ACP peptides on the basis of their amino acid fraction, global hydrophobicity, global hydrophobic movement, and global charge. Result suggests a significant difference in the frequency distribution of G, I, K, and L amino acids between two datasets. Additionally, ACP sequences depict a relatively higher positive global charge in compression to non-ACP that tends towards a relatively higher global negative charge. In addition to this, higher global hydrophobicity and hydrophobic movement has been observed in the ACP dataset. Interestingly, ACPs and non-ACPs show separation when plotted on the basis of their global hydrophobicity, global hydrophobic movement, and global change on a 3D scatter plot (Figure 4). Fisher’s test was used to explore the feature space expressed by hydropathy and the volume of amino acids. The example data set depicts significant over-representation of ACP sequences with larger hydrophobic amino acids. On the other hand, smaller hydrophilic residues are more frequent among sequences present in non-ACP groups (Figure 5).

**Figure 4.**
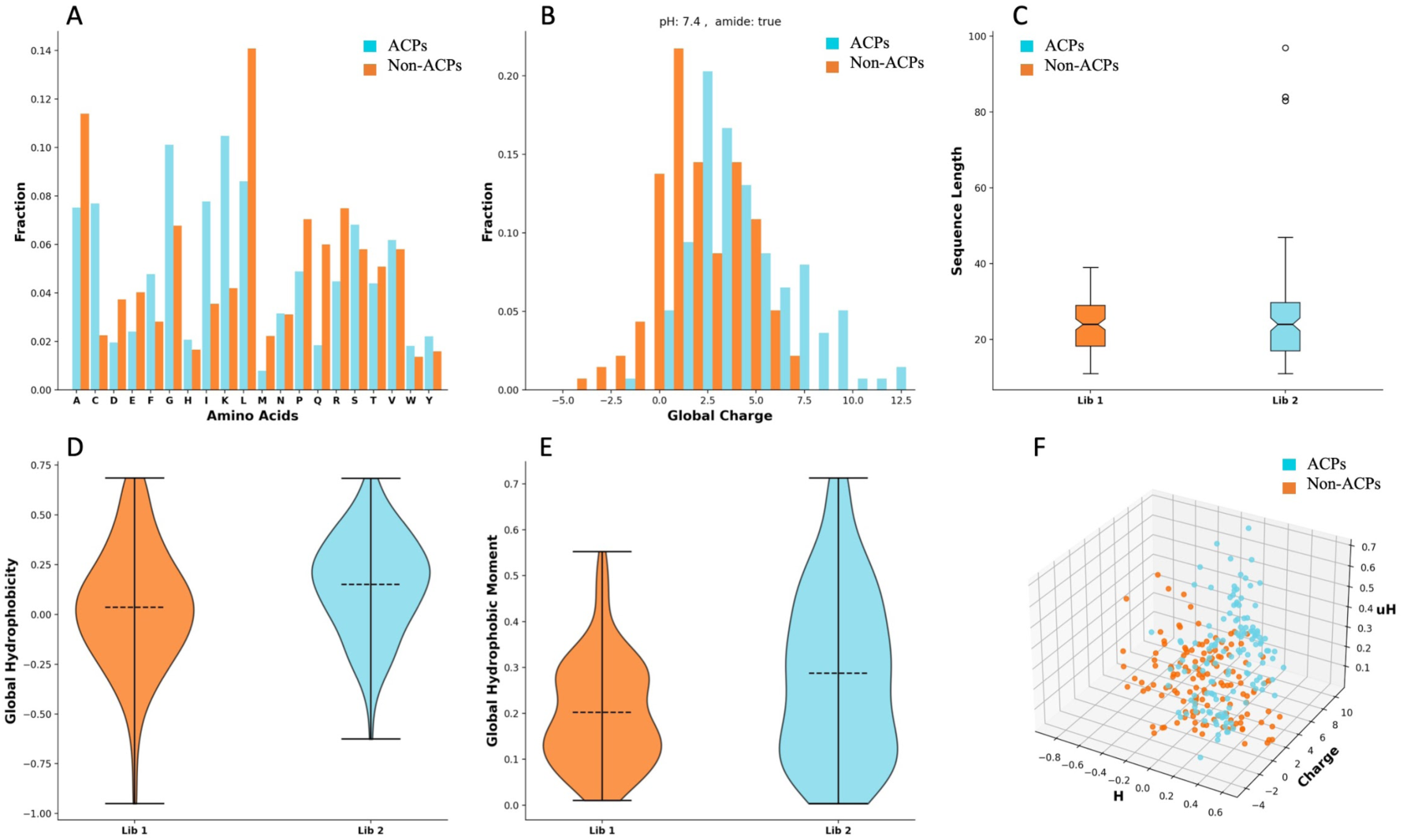
ACPs and non-ACPs datasets were compared and represented with a summary plot. A) Amino acid frequency distribution plot shows a significant difference in the frequency distribution of G, I, K, and L amino acids between ACPs and non-ACPs. B) Global charge distribution shows a higher positive charge among the ACPs, while overall higher negative charge occurs among non-ACPs sequences. C) There are no significant differences observed in the length distribution of ACPs and non-ACPs, except few outliers. D) ACPs and non-ACPs show differences in global hydrophobicity E) A relatively smaller hydrophobic moment has been observed in the non-ACPs in comparison to the ACPs. F) 3D scatter plot of global hydrophobicity, global hydrophobic movement and global charge showed separation between ACPs and non-ACPs.

**Figure 5.**
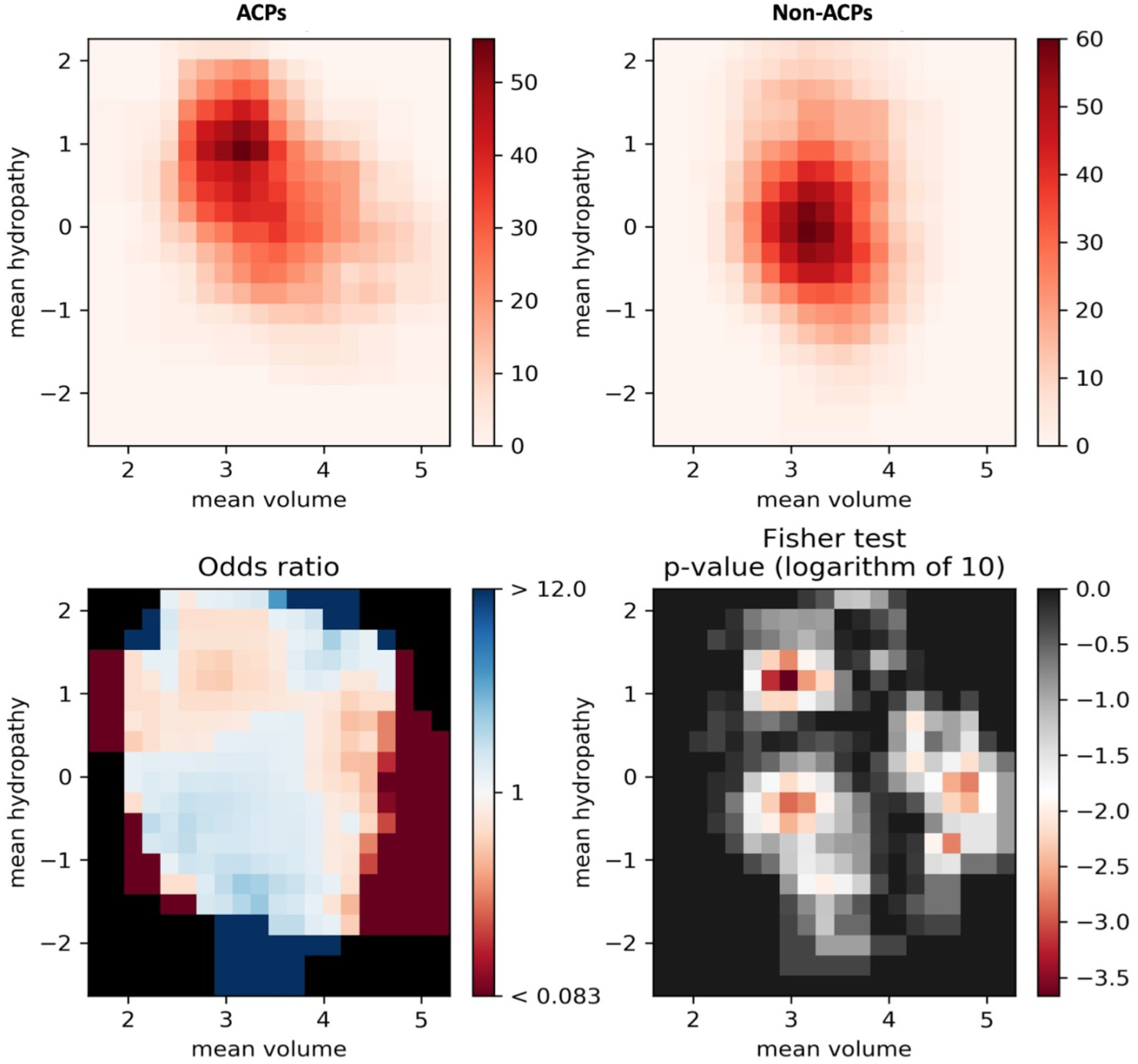
Feature space visualization of ACPs and non-ACPs. ACPs and non-ACPs in the feature space represented by their mean hydropathy and amino acid volume. The sequences with larger hydrophobic amino acids are more frequent in ACPs in comparison to non-ACPs.

### Example data results

Accuracy and the performance of all machine algorithms majorly depend upon how robustly a model has been trained, including the size and the complexity of the test dataset. Supervised classification methods have been commonly used to construct statistical models. Routinely used performance measures precision, recall, f1, measure, accuracy and AUC, as described above, have been incorporated to evaluate the performance of the enlisted machine learning algorithms. Results suggest that data normalization plays an important role in machine learning modeling and improves the performance of various machine learning algorithms. We observed that normalization significantly improves the performance of almost all the algorithms except RFC, which shows a negligible effect of normalization on machine learning models (Figure 6). The descriptor set is an important variable that affects the performance of ML algorithms and plays a crucial role. Here we examined the effect of different descriptors on an ML algorithm and vice versa. First, we compare the impact of descriptors on the ML performance, we found that ML algorithms exhibit comparatively higher performance when trained on CTD descriptors in comparison to the other two, Moran and Geary Autocorrelation descriptors. Additionally, a relatively higher positive effect of normalization was also observed if the model was trained on CTD descriptors in comparison to the other two descriptor sets. We also assessed the performance of various ML algorithms in combination with different descriptor sets. We found that CGDC is very sensitive to normalization and exhibits a significant improvement when trained on different descriptor sets. Conversely, as described earlier, RFC shows relatively less sensitivity to normalization and remains almost unaffected against all the descriptor set. The other three algorithms, GBC, SVM, and LRC showed relatively improved (Figure 6) performance after normalization in the case of all the three conventional descriptors. Interestingly, Word Vector descriptors, which is commonly used in natural language processing and adopted here to calculate text-based descriptors for machine learning modeling, outperformed models trained on other descriptors sets in order to classify peptide sequences on the basis of their biological properties. In addition to this, a drastic improvement has been observed in the performance of various algorithms after data normalization with models built upon Word Vector descriptors. We can clearly observe that almost all the algorithms, except SGDC, depict values close to 1.0 for all the accuracy measures. Results suggest that the majority of the algorithms show very high performance when trained on the Word Vector descriptor set in comparison to the other three descriptor sets (Figure 6). These results clearly indicate that the Word Vector descriptor outperformed the tested conventional descriptors and exhibits relatively high accuracy in machine learning model building. Workflows for each analysis, including ML modeling with the sequence-based descriptors (Supplementary Figure 1), ML modeling with word vector descriptors (Supplementary Figure 2), and peptide library analysis workflow (Supplementary Figure 3), are provided along with the example dataset as supplementary material.

**Figure 6.**
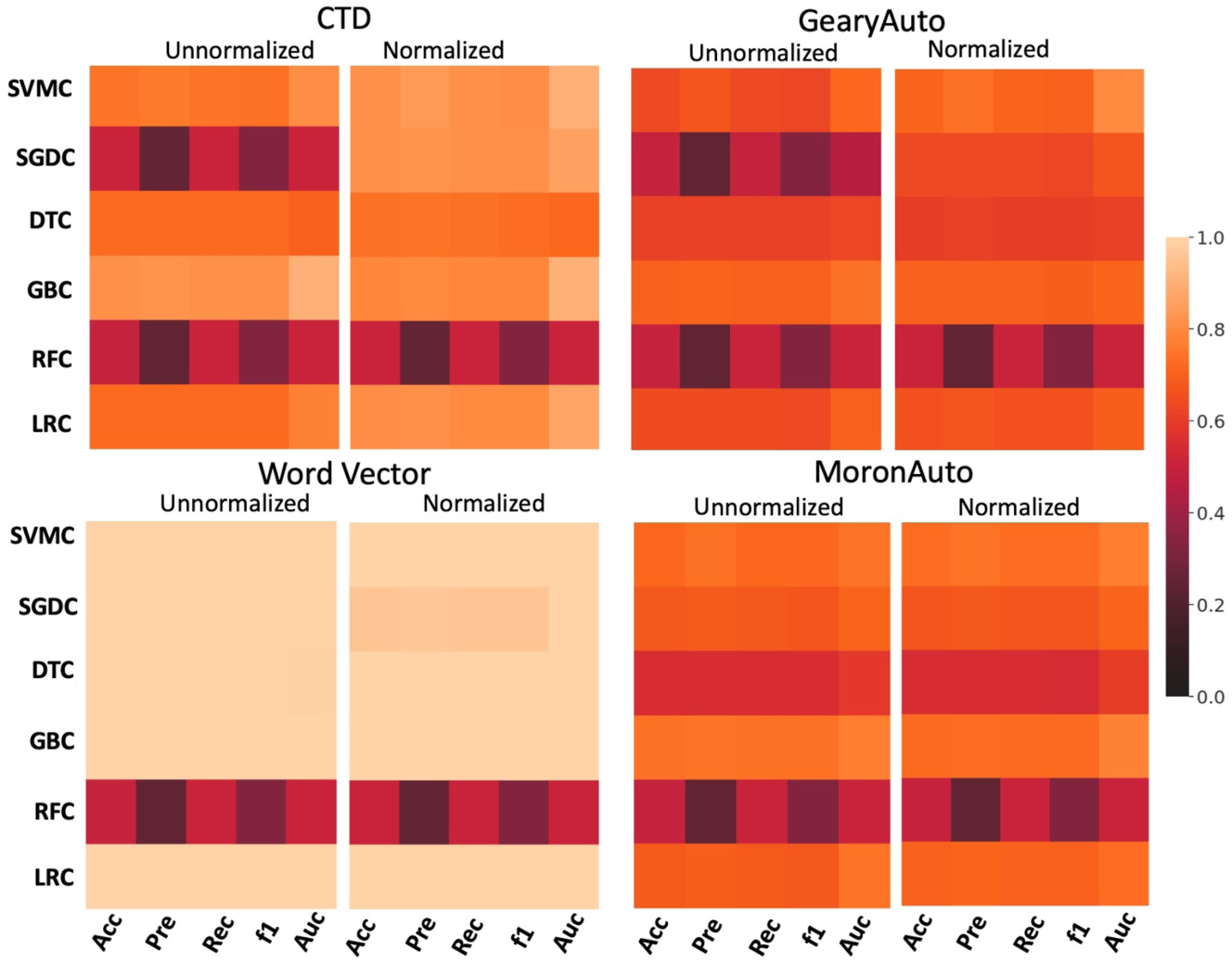
Assessment of the machine learning algorithms trained on four descriptor sets. Different performance measures accuracy, precision, recall, F1 score and mean AUC were calculated for six different algorithms with and without z-scaling normalization. Results suggest that the models trained on the word vector descriptors perform superior to the models trained on other descriptors.

## Conclusion

PDAUG leverages the Galaxy platform to provide a user friendly, reproducible peptide analysis environment. Researchers are able to assess the impact of differing tools, methods and algorithms, and to share and distribute their analysis results and workflows. This toolset provides researchers with access to GUI based tools for peptide library generation, feature analysis, data visualization and plotting, machine learning modeling, and dataset retrieval. PDAUG is released as open-source under the open source MIT license and with source code available from https://github.com/jaidevjoshi83/pdaug. Installation of PDAUG into a researcher’s Galaxy instance can be achieved using the point-and-click interface from the ToolShed. A docker image containing a PDAUG Galaxy system can also be obtained from https://hub.docker.com/r/jayadevjoshi12/galaxy_pdaug(https://github.com/jaidevjoshi83/docker_pdaug). Two interactive tutorials featuring this toolset, including workflows and sample datasets, combined with a detailed explanation of various tools, are available from https://training.galaxyproject.org/training-material/topics/proteomics/tutorials/peptide-library-data-analysis/tutorial.html and https://training.galaxyproject.org/training-material/topics/proteomics/tutorials/ml-modeling-of-anti-cancer-peptides/tutorial.html. A PDF version of these tutorials is also provided within the supplementary data.

## Supporting information

Tutorial - ML Modeling of Anti-cancer Peptides

Tutorial - Peptide Library Data Analysis

Galaxy Workflow Files

## Contributions

All authors contributed, read, and approved the manuscript.

## Competing interests

Daniel Blankenberg has a significant financial interest in GalaxyWorks, a company that may have a commercial interest in the results of this research and technology. This potential conflict of interest has been reviewed and is managed by the Cleveland Clinic.

## Notes

https://github.com/jaidevjoshi83/pdaug

https://hub.docker.com/r/jayadevjoshi12/galaxy_pdaug

https://github.com/jaidevjoshi83/docker_pdaug

https://training.galaxyproject.org/training-material/topics/proteomics/tutorials/peptide-library-data-analysis/tutorial.html

https://training.galaxyproject.org/training-material/topics/proteomics/tutorials/ml-modeling-of-anti-cancer-peptides/tutorial.html

