## Supplementary Figures for "PDAUG - a Galaxy based toolset for peptide library analysis, visualization, and machine learning modeling"

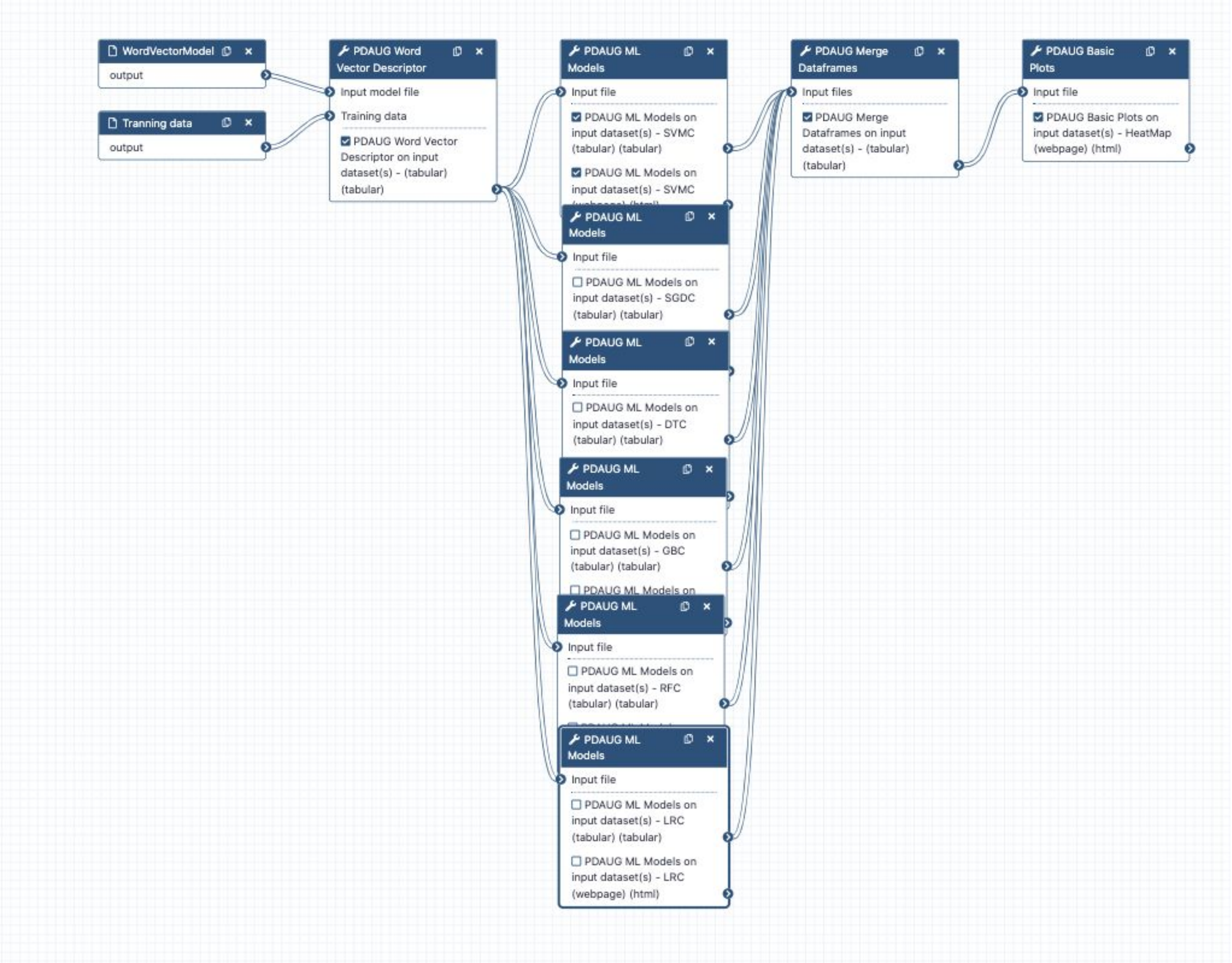


**Supplementary Figure 1**. Workflow to perform machine learning modeling based on word vector descriptors.

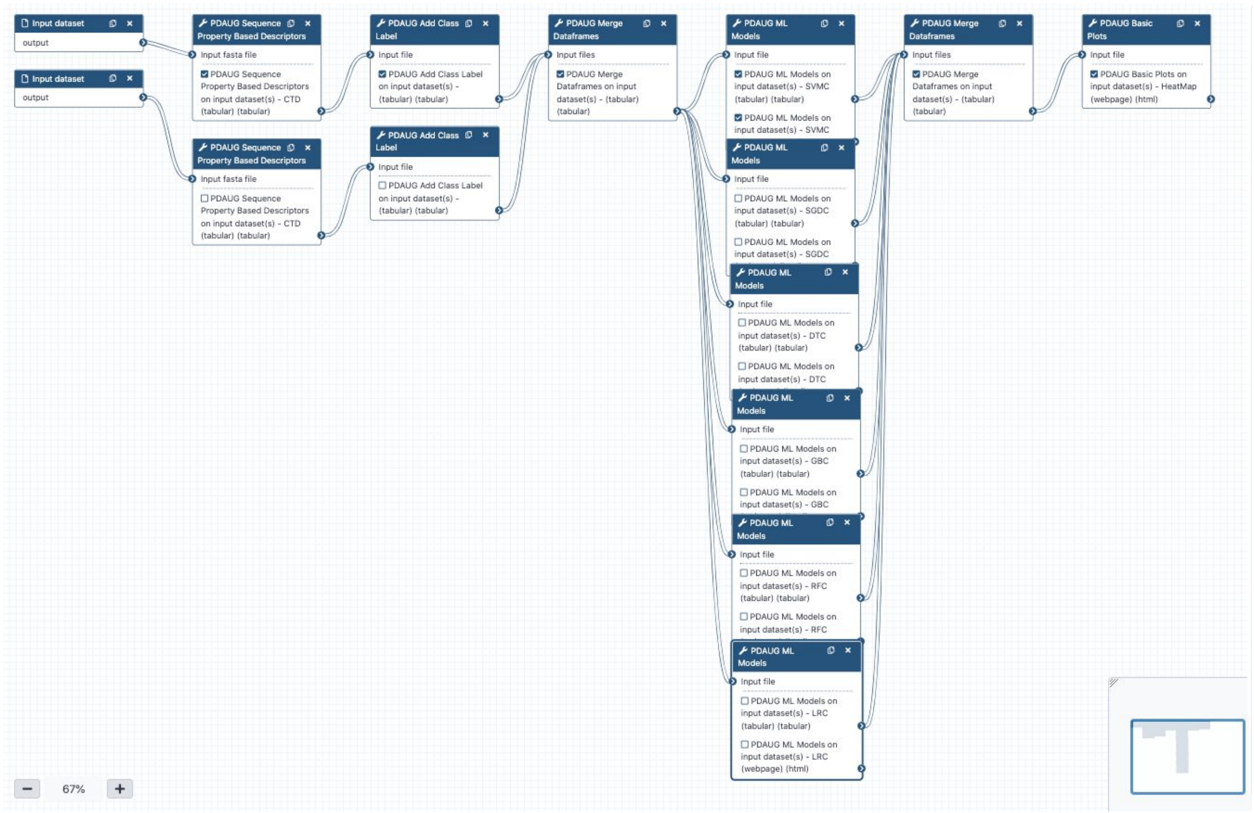


**Supplementary Figure 2.** Workflow to perform machine learning modeling based on CTD descriptors.

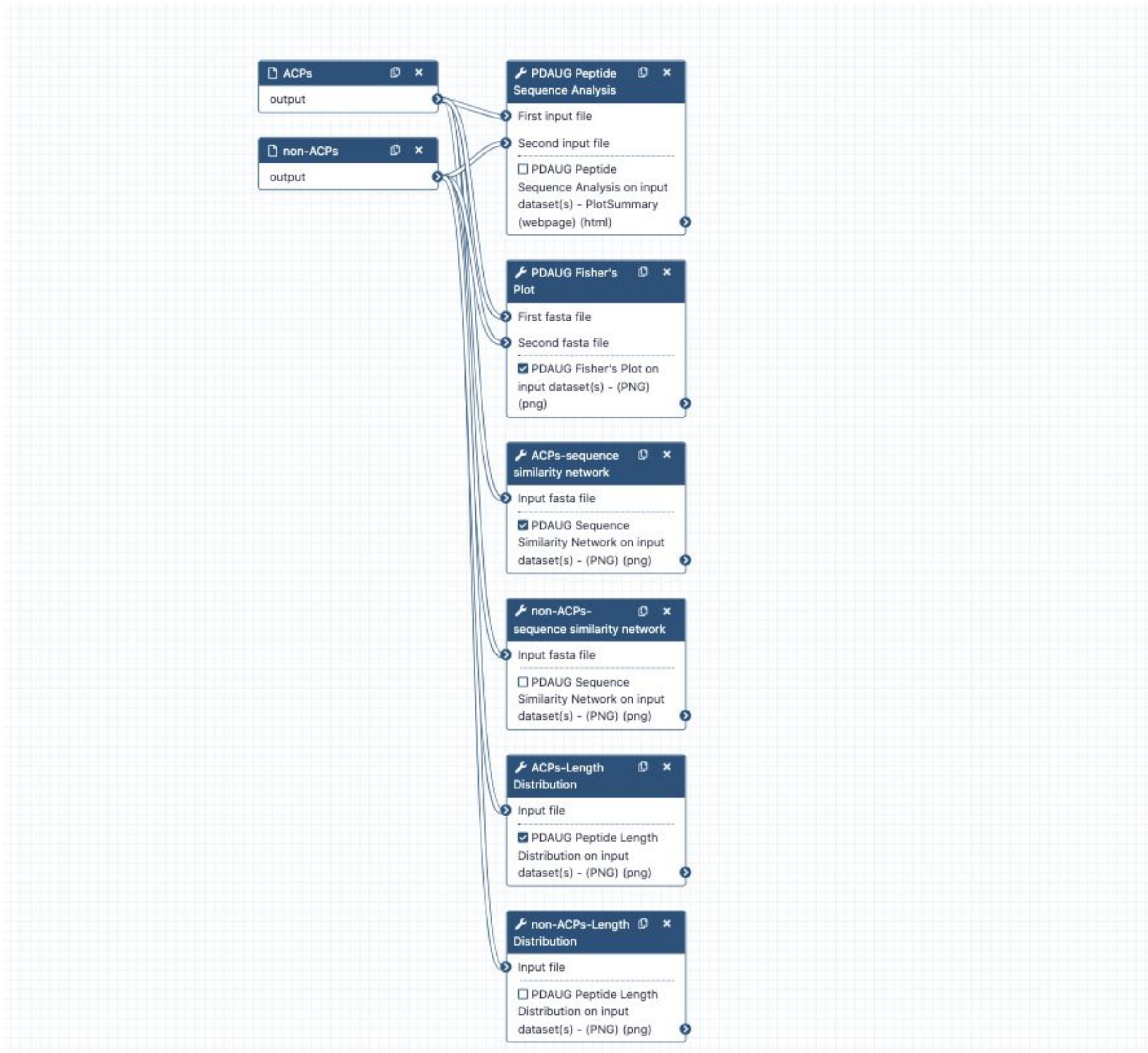


**Supplementary Figure 3**. Workflow to generate summary plot, Fisher's plot, sequence similarity network, and length distribution plot.
