## Supplementary material for "PDAUG - a Galaxy based toolset for peptide library analysis, visualization, and machine learning modeling": Tutorial - ML Modeling of Anti-cancer Peptides

### Machine Learning Modeling of Anticancer Peptides

By: Jayadev Joshi Daniel Blankenberg

Overview

**Questions**
- Which machine learning (ML) algorithm is superior in classifying anticancer peptides (ACPs) and non-anticancer peptides (non-ACPs)?

**Objectives**
- Learn, how to calculate peptide descriptor
- Learn, how to create training data set from features?
- Assessment of best ML algorithm in predicting anticancer peptide

**Requirements**
- Introduction to Galaxy Analyses

**Time estimation:** 30 minutes
**Level:** Intermediate
**Supporting Materials**
Datasets  Workflows  Available on these Galaxies
**Last modification:** Jan 22, 2021

#### Introduction

Biological molecules such as proteins, peptides, DNA, and RNA can be represented by their biochemical or sequence-based properties. These properties can be utilized to deduce biological meanings using ML modeling. A descriptor or feature is the quantitative or qualitative measure of a property that is associated with a sequence. For example, a chemical compound can be described via its charge chemical formula, molecular weight, number of rotatable bonds, etc. Similarly, several properties can be deduced from the biological sequence that can be utilized to describe a biological activity such as anticancer property. Properties associated with a group of peptide sequences such as overall charge, hydrophobicity profile, or k-mer composition can be utilized to build an ML model and this model can be used to predict biological properties of unknown peptides. Several computational methods have been proven very useful in the initial screening and prediction of peptides for various biological properties. These methods have emerged as effective alternatives to the lengthy and expensive traditional experimental approaches. Finding ACPs through wet-lab methods is costly and time-consuming; thus, the development of an efficient computational approach is useful to predict potential ACP peptides before wet-lab experimentation. In this tutorial, we will be discussing how peptide-based properties like charge, hydrophobicity, the composition of amino acids, etc. can be utilized to predict the biological properties of peptides. Additionally, we will learn how to use different utilities of the Peptide Design and Analysis Under Galaxy (PDAUG) package to calculate various peptide-based descriptors and use these descriptors for ML modeling. We will use CTD (composition, transition, and distribution) descriptor to define peptide sequences in the training set and will test 6 different ML algorithms. We will also assess the effect of normalization on the accuracy of ML models.

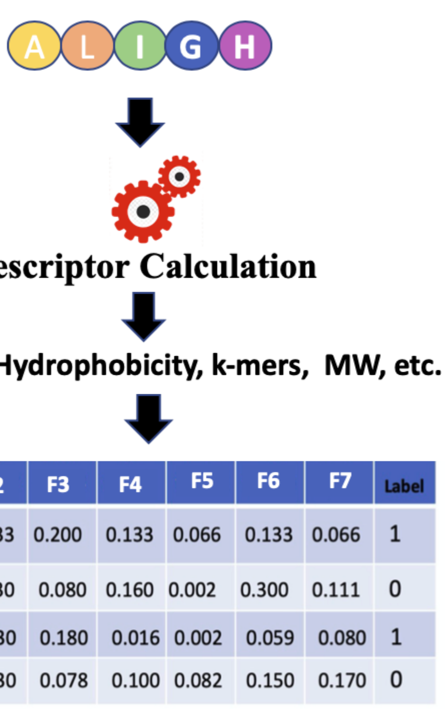


**Figure 1:** ML algorithms use numerical representation of a sequence-based properties for model building

In **Figure 1** features or descriptors are represented with (F1, F2, F3, etc.) and in binary classification, usually, class labels are represented by 0 or 1.

#### Easy access to tools, workflows and data from the docker image

An easy way to install and use the PDAUG toolset, and follow this tutorial is via a prebuilt docker image equipped with a PDAUG toolset, workflow, and data library. A prebuilds docker image can be downloaded and run by typing a simple command at the terminal after installing docker software on any operating system.

> Hands-on: Easy access of tools, workflows and data from docker image
>
> 1. Downloading the docker image from the docker hub using `docker pull jayadevjoshi12/galaxy_pdaug:latest` command.
> 2. Running the container with latest PDAUG tools `docker run -i -t -p 8080:80 jayadevjoshi12/galaxy_pdaug:latest` .
> 3. Workflow is available under the workflow section, use `admin` as username and `password` as a password to login as an administrator of your galaxy instance.
> 4. Use `admin` as username and `password` as a password to login galaxy instance, which is available at localhost to access workflow and data.

Agenda

In this tutorial, we will cover:
1. Training data set
2. Calculating Peptide Descriptors
3. Preparing a traning data set
4. Applying 6 different ML algorithms on the training data set
5. Results assessment

#### Training data set

A high-quality dataset was retrieved from a previously published work Hajisharifi *et al.* 2014. In the, ML balance training set (an equal number of positive and negative samples) is always recommended. However, an imbalance training set can also be handled, and sometimes, to assess the robustness of the model an imbalanced training set is intentionally introduced. The objective of this tutorial is to provide a basic introduction of ML in peptide research hence we will use a balanced training dataset with an equal number of positive (ACPs) and negative (non-ACPs) data. A simple python code was applied to randomly select and removed several non-ACPs, which reduces their number from 205 to 138. The final training set contains 138 ACPs and 138 non-ACPs. The length distribution of the positive dataset is somewhat different from the negative dataset. Peptide length is an important feature in determining biological activity. Based on their length we assess the differences between ACPs and non-ACPs, and we found that except for a few outliers both ACPs and non-ACPs show a mean length of 40 and 32 respectively.

##### Get data

> Hands-on: Data upload
>
> 1. Create a new history for this tutorial
> 2. Import the files from Zenodo or from the shared data library
>
> ```
> https://zenodo.org/record/4111092/files/ACPs.fasta
> https://zenodo.org/record/4111092/files/non_ACPs.fasta
> ```
>
> Tip: Importing data via links
>
> Tip: Importing data from a data library
>
> 3. Rename the datasets to their basename (ACPs.fasta, non_ACPs.fasta)
> 4. Check that the datatype is correctly set to fasta
>
> Tip: Changing the datatype

#### Calculating Peptide Descriptors

In this step we will calculate CTD descriptos. Composition describptors are defined as the number of amino acids of a particular property divided by total number of amino acids. Transition descriptors are representd as the number of transition from a particular property to different property divided by (total number of amino acids − 1). Distribution descriptors are derived by chain length and the amino acids of a particular property are located on this length Govindan and Nair 2013.

> Hands-on: Calculating CTD descriptors for ACPs and non-ACPs
>
> 1. **PDAUG Sequence Property Based Descriptors** with the following parameters:
>    - *"Input fasta file"*: `ACPs.fasta` (output of **Input dataset**)
>    - *"DesType"*: `CTD`
> 2. **PDAUG Sequence Property Based Descriptors** with the following parameters:
>    - *"Input fasta file"*: `non_ACPs.fasta` (output of **Input dataset**)
>    - *"DesType"*: `CTD`

#### Preparing a traning data set

We will combine the ACPs and non-ACPs data set as a single tabular data and will add the class label.

##### Adding class labels to the training data

In Binary classification, ML algorithms classify the elements of a set into two groups based on a classification rule. Binary classes are usually represented by `0` and `1` . In our example, we are finding peptides with anticancer properties, therefore we can denote all the peptides with anticancer properties as `1` and peptides with non-anticancer properties as `0` . In general, samples represented with `1` also describe as positive data, and samples with `0` labels as negative data. Data with multi-class classification problems can also be represented by `0, 1, 2, 3,` etc. In addition to this, in ML the class labels can also be represented by a string such as "anticancer" and "non-anticancer" or "treated" and "untreated". However, there are several ML tools and libraries prefer numerical class label over a string therefore, in this tutorial, `0` and `1` will be used as class labels.

> Hands-on: Adding class labels to the tabular data
>
> 1. **PDAUG Add Class Label** with the following parameters:
>    - *"Input file"*: `PDAUG Sequence Property Based Descriptors on data 1 - CTD (tabular)` (output of **Peptide Sequence Descriptors**)
>    - *"Class Label"*: `1`
> 2. **PDAUG Add Class Label** with the following parameters:
>    - *"Input file"*: `PDAUG Sequence Property Based Descriptors on data 2 - CTD (tabular)` (output of **Peptide Sequence Descriptors**)
>    - *"Class Label"*: `0`

##### Merging ACPs and non-ACPs samples to create a traning dataset

In previous steps, we labeled positive data or ACPs as "1", and negative data or non-ACPs as "0", now we can combine these two tabular datafiles as one training dataset. In this step, we will merge these tabular detasets as a labeled training data set.

> Hands-on: Merging two tabular files
>
> 1. **PDAUG Merge Dataframes** with the following parameters:
>    - *"Input files"*: `PDAUG Add Class Label on data 3 - (tabular)` (output of **Add Class Label**), `PDAUG Add Class Label on data 4 - (tabular)` (output of **Add Class Label**)

#### Applying 6 different ML algorithms on the training data set

In this step, we will apply six ML algorithms Linear Regression Classifier (LRC), Random Forest Classifier(RFC), Gaussian naive Bayes Classifier (GBC), Decision Tree Classifier (DTC), Stochastic Gradient Descent Classifier (SGDC) & Support Vector Machine Classifier (SVMC) with 10 fold cross-validation on the training data. In cross-validation, positive and negative data are randomly divided into 10 parts each set has the 10th part of active as well as inactive peptides. The algorithm was trained on the 9 sets and the prediction was made on the remaining 10th set. This process was repeated for every set. Thus the final performance scores are calculated as a mean of all the folds. We used min-max to normalize the data before ML modeling. The entire workflow was applied to the four descriptor sets and accuracy was estimated based on accuracy, precision, recall, f1, and AUC.

> Hands-on: Applying 6 ML algorithms on the training data set
>
> 1. **PDAUG ML Models** with the following parameters:
>    - *"Input file"*: `PDAUG Merge Dataframes on data 6 and data 5 - (tabular)` (output of **Merge dataframes**)
>    - *"Select Machine Learning algorithms"*: `LRC`
>      - *"Select advanced parameters"*: `No, use program defaults.`
>    - *"Choose the Test method"*: `Internal`
>    - *"Cross validation"*: `10`
> 2. **PDAUG ML Models** with the following parameters:
>    - *"Input file"*: `PDAUG Merge Dataframes on data 6 and data 5 - (tabular)` (output of **Merge dataframes**)
>    - *"Select Machine Learning algorithms"*: `RFC`
>      - *"Specify advanced parameters"*: `No, use program defaults.`
>    - *"Choose the Test method"*: `Internal`
>    - *"Cross validation"*: `10`
> 3. **PDAUG ML Models** with the following parameters:
>    - *"Input file"*: `PDAUG Merge Dataframes on data 6 and data 5 - (tabular)` (output of **Merge dataframes**)
>    - *"Select Machine Learning algorithms"*: `GBC`
>      - *"Specify advanced parameters"*: `No, use program defaults.`
>    - *"Choose the Test method"*: `Internal`
>    - *"Cross validation"*: `10`
> 4. **PDAUG ML Models** with the following parameters:
>    - *"Input file"*: `PDAUG Merge Dataframes on data 6 and data 5 - (tabular)` (output of **Merge dataframes**)
>    - *"Select Machine Learning algorithms"*: `DTC`
>      - *"Specify advanced parameters"*: `No, use program defaults.`
>    - *"Choose the Test method"*: `Internal`
>    - *"Cross validation"*: `10`
> 5. **PDAUG ML Models** with the following parameters:
>    - *"Input file"*: `PDAUG Merge Dataframes on data 6 and data 5 - (tabular)` (output of **Merge dataframes**)
>    - *"Select Machine Learning algorithms"*: `SGDC`
>      - *"Specify advanced parameters"*: `No, use program defaults.`
>    - *"Choose the Test method"*: `Internal`
>    - *"Cross validation"*: `10`
> 6. **PDAUG ML Models** with the following parameters:
>    - *"Input file"*: `PDAUG Merge Dataframes on data 6 and data 5 - (tabular)` (output of **Merge dataframes**)
>    - *"Select Machine Learning algorithms"*: `SVMC`
>      - *"Specify advanced parameters"*: `No, use program defaults.`
>    - *"Choose the Test method"*: `Internal`
>    - *"Cross validation"*: `10`

#### Results assessment

##### Merging results in one file

In previous steps we have trained the ML models, these models return a TSV that captures performance measures of these algorithms. We used the `Merge Data Frame` tool to combine these results as one file in this step.

> Hands-on: Merging result as one tabular file
>
> 1. **PDAUG Merge Dataframes** with the following parameters:
>    - *"Input files"*: `PDAUG ML Models on data 7 - LRC (tabular)` (output of **ML Models**), `PDAUG ML Models on data 7 - RFC (tabular)` (output of **ML Models**), `PDAUG ML Models on data 7 - GBC (tabular)` (output of **ML Models**), `PDAUG ML Models on data 7 - DTC (tabular)` (output of **ML Models**), `PDAUG ML Models on data 7 - SGDC (tabular)` (output of **ML Models**), `PDAUG ML Models on data 7 - SVMC (tabular)` (output of **ML Models**)

##### Creating a final heat map to assess the results

In the final step, a heat map will be generated which represents performance measures of various algorithms. We applied five different performance measures, accuracy, recall, F1-score, precision, and mean AUC (Area Under Curve) score.

> Hands-on: Plotting the results
>
> 1. **PDAUG Basic Plots** with the following parameters:
>    - *"Data plotting method"*: `Heat Map`
>      - *"Input file"*: `PDAUG Merge Dataframes on data 18, data 16, and others - (tabular)` (output of **PDAUG Merge Dataframes**)
>      - *"Index Column"*: `Algo`
>      - *"Label for x-axis"*: `Performance Measures`
>      - *"Label for y-axis"*: `ML algorithms`

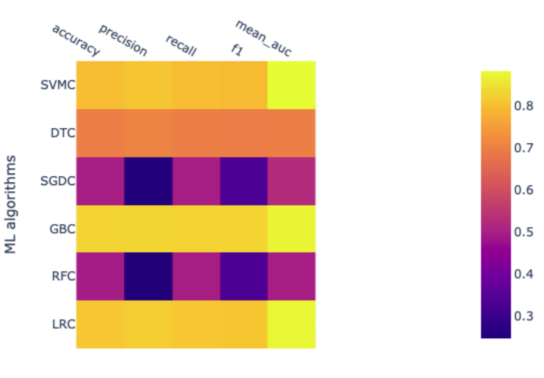


**Figure 2:** Heatmap represents the performance of 6 ML algorithms

The performance of ML algorithms can be assessed by commonly used performance measures represented in **Figure 2**.

- **Accuracy** is described as correctly predicted instances and calculated based on true positive (TP) and true negative (TN) divided by TP, TN, false positive (FP), and false-negative (FN).
- **AUC** is Area under ROC curve, where ROC is a receiver operating characteristic. AUC represents the area covered by ROC.
- **F1** measures also an important estimate in model accuracy and can be defined as a harmonic mean of precision.
- **Precision** also known as the probability of positive values (PPV), is summarised as the probability of currently predicted positive instances and estimated based on TP and FP.
- **Recall** also known as sensitivity, is defined as the estimation of the percentage of the correctly predicted positive instances and is also calculated with TP and FP.

The value for each of these estimates falls between 0 and 1, and larger values indicating a better performance and accuracy. The brighter yellow color shows high-performance while the blue color shows a lower score. Heat map suggests that algorithms GBC, LRC, and SVMC show high performs in comparison to the other three. Better performance means these classifiers have been able to classify ACPs and non-ACPs more accurately than others. DTC shows an intermediate performance while RFC and SGDC performed poorly on this data set. Finally, we learn how to calculate features, how to utilize these features to build ML models, and how we can assess the performance of an ML model. In the future, advanced parameters of this algorithm can be assessed to improve the performance of the models. Additionally, several features can be calculated other than CTD and utilized to build ML models and assess performance.

#### Conclusion

In this tutorial, we learn how to utilize the quantitative properties of peptide sequences and apply the ML algorithms to predict the biological properties of the peptide sequence.

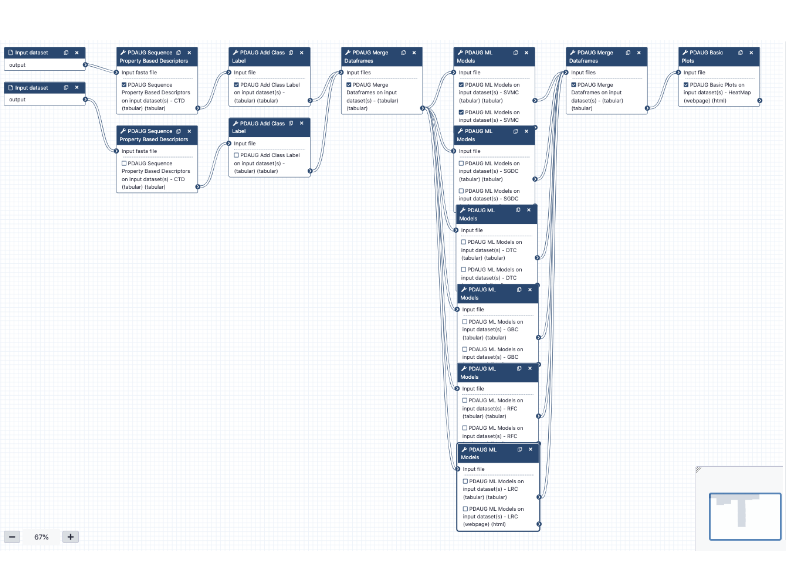

**Figure 3:** Workflow

#### Useful literature

Further information, including links to documentation and original publications, regarding the tools, analysis techniques and the interpretation of results described in this tutorial can be found here.


#### Feedback

Did you use this material as an instructor? Feel free to give us feedback on how it went.

Help us improve this content!

Your feedback helps us improve this tutorial and will be considered in future revisions.

This feedback should be ONLY ABOUT THE MANUAL; if you encountered problems with the Galaxy server or if tools were missing, please contact the administrators of the Galaxy server you were using.

We do not store any personal identifying information.

How much did you like this tutorial?

1 2 3 4 5

#### Citing this Tutorial


1. Jayadev Joshi, Daniel Blankenberg, 2021 **Machine Learning Modeling of Anticancer Peptides (Galaxy Training Materials)**. /training-material/topics/proteomics/tutorials/ml-modeling-of-anti-cancer-peptides/tutorial.html Online; accessed Mon Feb 01 2021
2. Batut et al., 2018 **Community-Driven Data Analysis Training for Biology** Cell Systems 10.1016/j.cels.2018.05.012


BibTeX

##### Congratulations on successfully completing this tutorial!

This material is the result of a collaborative work. Thanks to the Galaxy Training Network and all the contributors (Jayadev Joshi, Daniel Blankenberg)!

Found a typo? Something is wrong in this tutorial? Edit it on GitHub.

The content of the tutorials and website is licensed under the Creative Commons Attribution 4.0 International License.
