## Supplementary material for "PDAUG - a Galaxy based toolset for peptide library analysis, visualization, and machine learning modeling": Tutorial - Peptide Library Data Analysis

Overview

**Questions**
- How to utilize quantitative properties of amino acids and peptide sequence to analyse peptide data?

**Objectives**
- Calculate descriptors
- Quantitative analysis of peptide sequence properties

**Requirements**
- Introduction to Galaxy Analyses

**Time estimation:** 20 minutes
**Level:** Intermediate
**Supporting Materials**
Workflows Available on these Galaxies
**Last modification:** Jan 22, 2021

### Introduction

Several computational methods have been proven very useful in the initial screening and prediction of peptides for various biological properties. These methods have emerged as effective alternatives to the lengthy and expensive traditional experimental approaches. Properties associated with a group of peptide sequences such as overall charge, hydrophobicity profile, or k-mer composition can be utilized to compare peptide sequences and libraries. In this tutorial, we will be discussing how peptide-based properties like charge, hydrophobicity, the composition of amino acids, etc. can be utilized to analyze the biological properties of peptides. Additionally, we will learn how to use different utilities of the Peptide Design and Analysis Under Galaxy (PDAUG) package to calculate various peptide-based descriptors, and use these descriptors and feature spaces to build informative plots.

#### Easy access to tools, workflows and data from the docker image

An easy way to install and use the PDAUG toolset, and follow this tutorial is via a prebuilt docker image equipped with a PDAUG toolset, workflow, and data library. A prebuilds docker image can be downloaded and run by typing a simple command at the terminal after installing docker software on any operating system.

Hands-on: Easy access of tools, workflows and data from docker image

1. Downloading the docker image from the docker hub using `docker pull jayadevjoshi12/galaxy_pdaug:latest` command.
2. Running the container with latest PDAUG tools `docker run -i -t -p 8080:80 jayadevjoshi12/galaxy_pdaug:latest`.
3. Workflow is available under the workflow section, use `admin` as username and `password` as a password to login as an administrator of your galaxy instance.
4. Use `admin` as username and `password` as a password to login galaxy instance, which is available at localhost to access workflow and data.

Agenda

In this tutorial, we will cover:
1. Peptide Data
2. Converting tabular data into fasta format
3. Analyzing peptide libraries (AMPs and TMPs) based on features and feature space
4. Assessing the relation between peptide features by 3D scatter plot

### Peptide Data

Several inbuilt data sets have been provided with the tool `PDAUG Peptide Data Access`. The antimicrobial peptides (AMPs) versus transmembrane peptides (TMPs) dataset was used as an example data set to understand the overall relation between features and biological properties of peptides. AMPs consist of an intersection of all activity annotations of the APD2 and CAMP databases, where gram-positive, gram-negative, and antifungal exact matches were observed. TMPs were extracted from alpha-helical transmembrane regions of proteins for classification.

Hands-on: Fetching inbuild data

1. **PDAUG Peptide Data Access** with the following parameters:
   - *"Datasets"*: `AMPvsTMP`

### Converting tabular data into fasta format

`PDAUG Peptide Data Access` tool returns data as a tabular file that contains sequences from both the classes. In order to utilize this data in the next steps, first we need to convert tabular data into fasta format. If data contains sequences from two different classes `PDAUG TSVtoFASTA` tool converts and splits data into two separate files for each of the class, AMPs, and TMPs. The reason behind converting and splitting the data is that all the downstream tools require two separate files if we are comparing two different peptide classes or calculating features.

Hands-on: Converting tabular data into fasta formate

1. **PDAUG TSVtoFASTA** with the following parameters:
   - *"Input file"*: `PDAUG Peptide Data Access - AMPvsTMP (tabular)` (output of **PDAUG Peptide Data Access**)
   - *"Peptide Column"*: `name`
   - *"Method to convert data"*: `Split Data By Class Label`
   - *"Column with the class label"*: `class label`

### Analyzing peptide libraries (AMPs and TMPs) based on features and feature space

#### Summary Plot for peptide libraries

In this step, we utilize `PDAUG Peptide Sequence Analysis` tool to compare peptide sequences based on hydrophobicity, hydrophobic movement, charge, amino acid fraction, and sequence length and create a summary plot.

Hands-on: Generating a summary plot to assess peptide dataset

1. **PDAUG Peptide Sequence Analysis** with the following parameters:
   - *"Analysis options"*: `Plot Summary`
     - *"First input file"*: `PDAUG TSVtoFASTA on data 1 - first (fasta)` (first output of **PDAUG TSVtoFASTA**)
     - *"Second input file"*: `PDAUG TSVtoFASTA on data 1 - second (fasta)` (second output of **PDAUG TSVtoFASTA**)
     - *"first input file"*: `TMPs`
     - *"Second input file"*: `AMPs`

Questions

What can be concluded from the summary plot based on different properties?

Solution

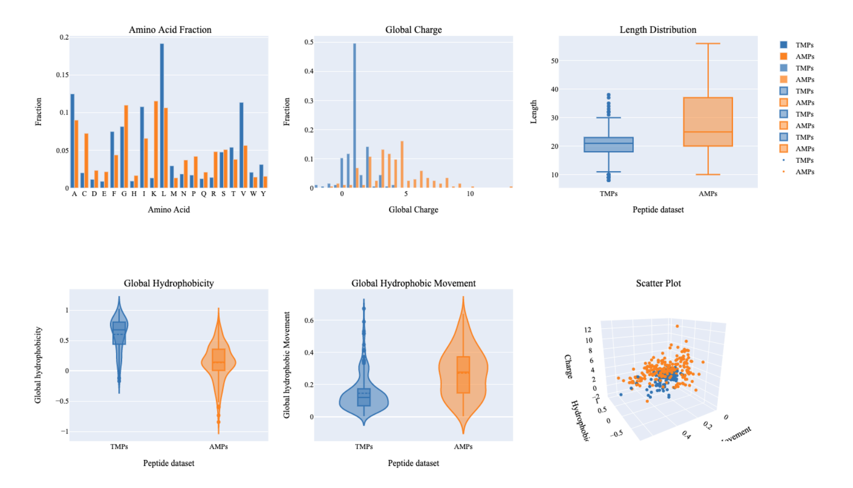


**Figure 1:** Summary plot shows comparison between AMPs and TMPs

#### Assessing feature space distribution

In this tool, we have used `PDAUG Fisher's Plot` that compares two peptide libraries based on the feature space using the Fisher test.

Hands-on: Generating a Fisher's plot to assess peptide dataset

1. **PDAUG Fisher's Plot** with the following parameters:
   - *"First fasta file"*: `PDAUG TSVtoFASTA on data 1 - first (fasta)` (first output of **PDAUG TSVtoFASTA**)
   - *"Second fasta file"*: `PDAUG TSVtoFASTA on data 1 - second (fasta)` (second output of **PDAUG TSVtoFASTA**)
   - *"Label for first population"*: `TMPs`
   - *"Label for second population"*: `AMPs`

Questions

What does Fisher's plot represents?

Solution

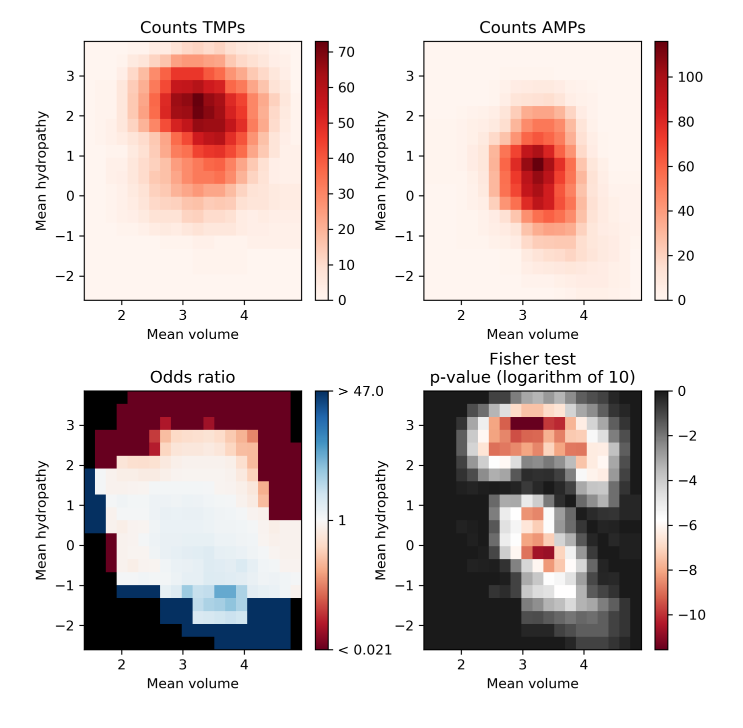


**Figure 2:** TMPs peptides show amino acids with larger hydrophobic residues in compare to AMPs

The AMPs and TMPs in the feature space represented by their mean hydropathy and amino acid volume. Fisher's plot shows that the sequences with larger hydrophobic amino acids are more frequent in TMPs in comparison to AMPs.

### Assessing the relation between peptide features by 3D scatter plot

#### Calculating Sequence Property-Based Descriptors

In this step we will calculate Composition, Transition and Distribution (CTD) descriptos. Composition describptors are defined as the number of amino acids of a particular property divided by total number of amino acids. Transition descriptors are representd as the number of transition from a particular property to different property divided by (total number of amino acids − 1). Distribution descriptors are derived by chain length and the amino acids of a particular property located on this length Govindan and Nair 2013.

Hands-on: Calculating descriptors for the peptide dataset

1. **PDAUG Sequence Property Based Descriptors** with the following parameters:
   - *"Input fasta file"*: `PDAUG TSVtoFASTA on data 1 - first (fasta)` (first output of **PDAUG TSVtoFASTA**)
   - *"Descriptor Type"*: `CTD`
2. **PDAUG Sequence Property Based Descriptors** with the following parameters:
   - *"Input fasta file"*: `PDAUG TSVtoFASTA on data 1 - second (fasta)` (second output of **PDAUG TSVtoFASTA** )
   - *"Descriptor Type"*: `CTD`

#### Adding the Class Label in both AMPs and TMPs

Class labels or target labels usually represents the class of peptides. Here in our data set, we have peptides, either as AMP or TMP. Since we have two classes we can represent these two classes with their actual labels AMPs and TMPs.

- **Adding Class Label (target labels) in AMPs and TMPs data**

Hands-on: Adding Class Labels (target labels) to the tabular data

1. **PDAUG Add Class Label** with the following parameters:
   - *"Input file"*: `PDAUG Sequence Property Based Descriptors on data 2 - CTD (tabular)` (output of **PDAUG Sequence Property Based Descriptors**)
   - *"Class Label"*: `TMPs`
2. **PDAUG Add Class Label** with the following parameters:
   - *"Input file"*: `PDAUG Sequence Property Based Descriptors on data 3 - CTD (tabular)` (output of **PDAUG Sequence Property Based Descriptors**)
   - *"Class Label"*: `AMPs`

#### Merging the two tabular data files

We utilize `PDAUG Merge Dataframes` to merge two tabular data files.

Hands-on: Merging two tabular data files

1. **PDAUG Merge Dataframes** with the following parameters:
   - *"Input files"*: `PDAUG Add Class Label on data 6 - (tabular)` (output of **PDAUG Add Class Label**), `PDAUG Add Class Label on data 7 - (tabular)` (output of **PDAUG Add Class Label**)
   - *"Option to merg data"*: `Merge data without adding class label`

#### Plotting CTD descriptor data as Scatter plot

Tool `PDAUG Basic Plots` will be used to compare two peptide libraries based on three CTD descriptors `SecondaryStrD1100`, `SolventAccessibilityD2001`, and `NormalizedVDWVD3050` respectively. A 3D scatter plot will be generated.

Hands-on: Generating a scatter plot to assess features

1. **PDAUG Basic Plots** with the following parameters:
   - *"Data plotting method"*: `Scatter Plot`
     - *"Input file"*: `PDAUG Merge Dataframes on data 9 and data 8 - (tabular)` (output of **PDAUG Merge Dataframes**)
     - *"Scatter Plot type"*: `3D`
       - *"First feature"*: `_SecondaryStrD1100`
       - *"Second feature"*: `_SolventAccessibilityD2001`
       - *"Third feature"*: `_NormalizedVDWVD3050`
     - *"Class label column"*: `Class_label`

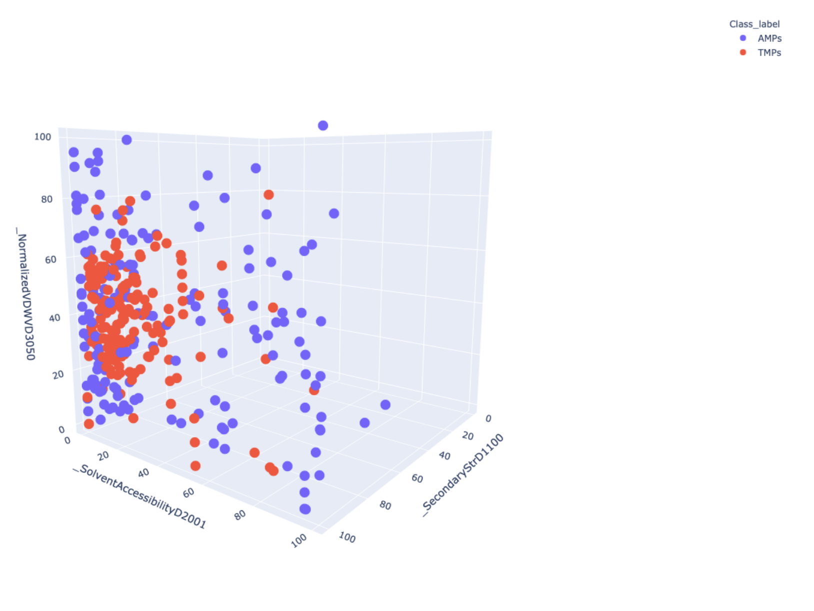


**Figure 3:** 3D scatter Plot shows relation between featues

**Figure 3** Represent 3D scattered plot generated based on the CTD descriptors. Red dots represent TMPs and blue dots represent AMPs. Based on these 3 features, we can observe that both groups do not show any clear separation or cluster in the 3D space.

In this tutorial, we learned how to utilize inbuild data, calculate features, and utilize descriptors or features to assess biological properties. We also learned how to utilize various utilities of `PDAUG` to generate useful plots to include in our peptide research.

### Conclusion

In this tutorial, we learned an example flexible and extensible analysis of peptide data using PDAUG tools. We generated various plots based on the quantitative properties of amino acids and peptide sequences.

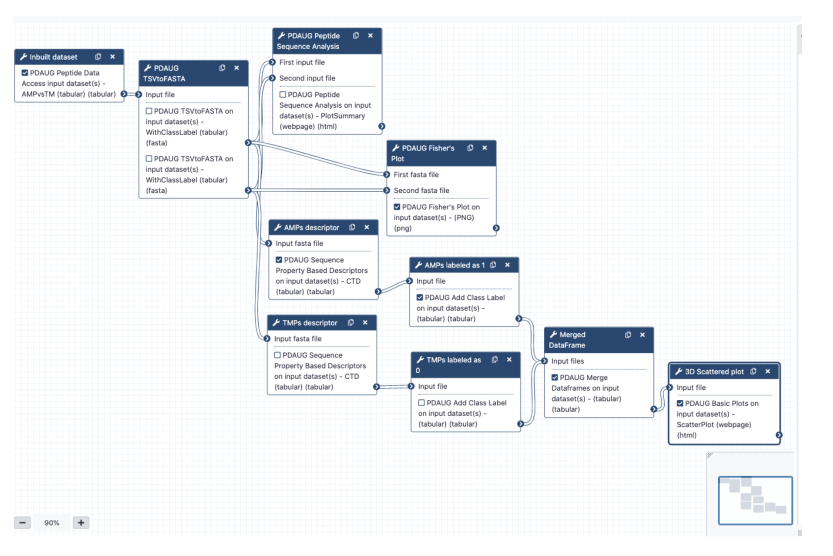


**Figure 4:** Workflow used

### Useful literature

Further information, including links to documentation and original publications, regarding the tools, analysis techniques and the interpretation of results described in this tutorial can be found here.

### Feedback

Did you use this material as an instructor? Feel free to give us feedback on how it went.

#### Help us improve this content!

Your feedback helps us improve this tutorial and will be considered in future revisions.

This feedback should be ONLY ABOUT THE MANUAL; if you encountered problems with the Galaxy server or if tools were missing, please contact the administrators of the Galaxy server you were using.

We do not store any personal identifying information.

How much did you like this tutorial?

1 2 3 4 5

### Citing this Tutorial

1. Jayadev Joshi, Daniel Blankenberg, 2021 **Peptide Library Data Analysis (Galaxy Training Materials)**. /training-material/topics/proteomics/tutorials/peptide-library-data-analysis/tutorial.html Online; accessed Mon Feb 01 2021
2. Batut et al., 2018 **Community-Driven Data Analysis Training for Biology** Cell Systems 10.1016/j.cels.2018.05.012

BibTeX

#### Congratulations on successfully completing this tutorial!

This material is the result of a collaborative work. Thanks to the Galaxy Training Network and all the contributors (Jayadev Joshi, Daniel Blankenberg)!

Found a typo? Something is wrong in this tutorial? Edit it on GitHub.

The content of the tutorials and website is licensed under the Creative Commons Attribution 4.0 International License.
